# scGEAToolbox: a Matlab toolbox for single-cell RNA sequencing data analysis

**DOI:** 10.1101/544163

**Authors:** James J. Cai

## Abstract

**Motivation:** Single-cell RNA sequencing (scRNA-seq) technology has revolutionized the way research is done in biomedical sciences. It provides an unprecedented level of resolution across individual cells for studying cell heterogeneity and gene expression variability. Analyzing scRNA-seq data is challenging though, due to the sparsity and high dimensionality of the data.

**Results:** I developed scGEAToolbox—a Matlab toolbox for scRNA-seq data analysis. It contains a comprehensive set of functions for data normalization, feature selection, batch correction, imputation, cell clustering, trajectory/pseudotime analysis, and network construction, which can be combined and integrated to building custom workflow. While most of the functions are implemented in native Matlab, wrapper functions are provided to allow users to call the “third-party” tools developed in Matlab or other languages. Furthermore, scGEAToolbox is equipped with sophisticated graphical user interfaces (GUIs) generated with App Designer, making it an easy-to-use application for quick data processing.

**Availability:** https://github.com/jamesjcai/scGEAToolbox

## 1 Introduction

Single-cell technologies, especially single-cell RNA sequencing (scRNA-seq), have revolutionized the way biologists and geneticists study cell heterogeneity and gene expression variability. Analyzing scRNA-seq data, however, is a challenging task due to the sparsity and dimensionality of the data. The sparsity problem is rooted from the limitation in the sensitivity of single-cell assay system; scRNA-seq data sets are often confounded by nuisance technical effects. The analyses of scRNA-seq data involve, in general, data filtering, normalization, feature selection, cell clustering, marker gene identification, cell type identification, pseudotime or trajectory analysis, gene regulatory network construction, and so on. For each of these analyses, there have many software tools for fulfilling the task. The majority of these tools are developed in computer languages such as R and python; few tools [e.g., SIMLR (Wang, et al., 2018) and SoptSC (Wang, et al., 2019)] are developed in Matlab, which is a scientific programming language providing strong mathematical and numerical support for the implementation of advanced algorithms. The basic data element of Matlab is the matrix; mathematical operations that work on arrays or matrices are built-in to the Matlab environment. Matlab comes with many toolboxes, such as statistics, bioinformatics, optimization, and image processing. Given scRNA-seq data is increasing exponentially over time, a new Matlab toolbox for such data analyses is highly desired.

## 2 Methods

I developed scGEAToolbox using Matlab v9.5 (R2018b). Functions were written in native Matlab and the app GUIs were created using the App Designer. The majority of functions accept two input arguments: `X` and `genelist`, where `X` is a *n×m* matrix holding expression values (e.g., UMIs) of *n* genes in *m* cells; `genelist` is an *n*×1 string array holding the names of genes. Main categories of functions of scGEAToolbox include: file input and output, data normalization, gene and cell filtration, detection of highly variable genes (HVGs), batch effect correction, dimensionality reduction, data visualization, cell clustering, trajectory analysis, and network construction. For each category, multiple algorithms were implemented. For example, for data normalization, two functions, `norm_libsize` and `norm_deseq`, were developed for the task of normalizing `X` using the two different methods, respectively. Furthermore, an “entry” function called `sc_norm` was developed to allow users to access the two functions using syntax like: `sc_norm`(`X, ‘type’, ‘libsize’`) and `sc_norm`(`X, ‘type’, ‘deseq’`). Accord-i ngly, the functionSignatures.json file was edited to specify the usage of all entry functions. The main GUI application in scGEAToolbox is called scGEApp. It contains the main panel with multiple tabs, namely *Load Data, Filter, Normalization, Batch Correction, Imputation, Feature Selection, Visualization, Clustering, Pseudotime*, and *Network.* On each tab panel, there are buttons for executing corresponding functions. For example, function for selecting cells by library size and selecting genes by the number of mapped reads are under *Filter*; functions for HVG selection are under *Feature Selection*; functions for t-SNE and PHATE are under *Visualization*. Under the main panel is a panel for viewing data matrices and result tables, where data and results can be exported into the workspace as variables or saved into external files. To expand its functionality, scGEAToolbox incorporates functions from several Matlab-based tools such as ComBat, MAGIC, McImpute, SIMLR, SinNLRR, SoptSC, PHATE, scDiffMap and GENIE3. In scGEAToolbox, these external tools can be accessed through corresponding wrapper functions such as `run_magic, run_simlr` and `run_genie3`. Furthermore, scGEAToolbox also includes wrapper functions for selected R functions such as UMAP, SCODE and Monocle.

**Fig. 1.**
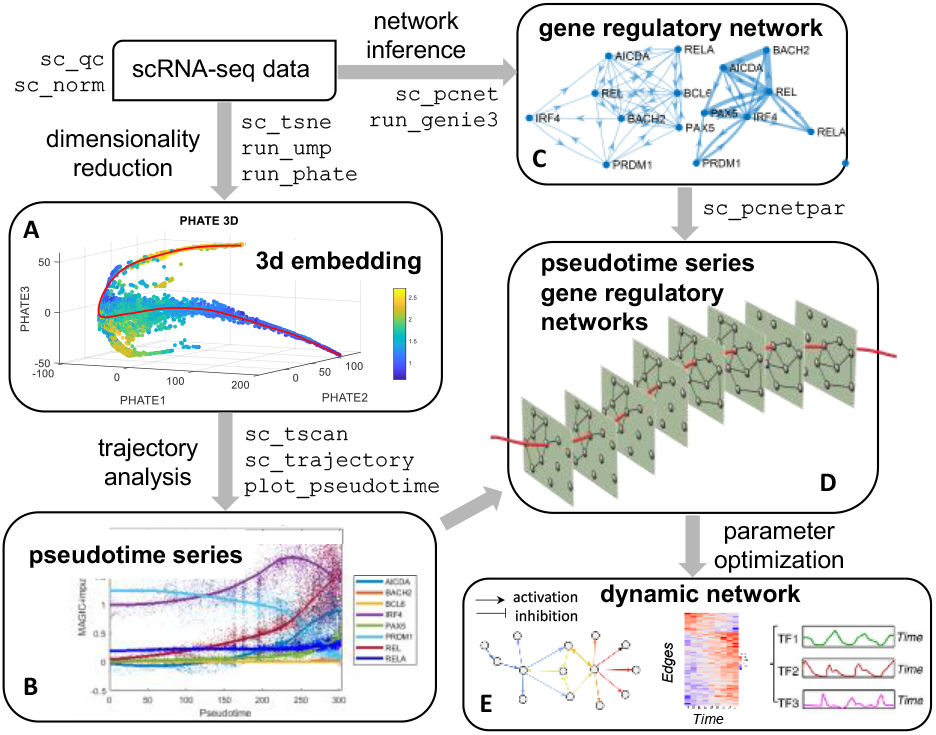
A software workflow built with scGEAToolbox for single-cell gene regulatory network (scGRN) analyses. High-dimensional scRNA-seq data is filtered, normalized, and used as input for two paths. The first is a combination of **(A)** dimensionality reduction and **(B)** trajectory/psedotime analysis to provide pseudotime-series data. The second is using network inference algorithms to generate **(C)** a global, coarse GRN structure. The integration of results from the two paths produces **(D)** pseudotime-series scGRNs, which can be further analyzed through regulatory modeling using parameter estimation algorithms to infer **(E)** a refined dynamic scGRN.

## 3 Results

scGEAToolbox has been developed to facilitate the analyses of scRNA-seq data in Matlab environment. It contains a comprehensive set of high-level functions, all taking the same input arguments: `X` and `genelist`. Such a uniform functional interface simplifies the syntax of function call, making scGEAToolbox an ideal algorithm prototyping tool. These high-level functions include, for example, `sc_hvg` and `sc_veg` for HVG detection (Brennecke, et al., 2013; Chen, et al., 2016), `sc_sc3` for cell clustering (Kiselev, et al., 2017), `sc_tscan` for trajectory analysis (Ji and Ji, 2016), and `sc_pcnet` for constructing gene regulatory network (GRN) (Gill, et al., 2010). The value of scGEAToolbox lies on the fact that its comprehensive sets of functions can be “recombined” in different ways to create new analytical workflows, providing customized solutions for real-world applications. **Fig. 1** shows an example of such workflows. This workflow is build using several scGEAToolbox functions and allows users to construct a series of gene regulatory networks (GRNs) from cells subsampled according to their pseudotime. The output of the workflow is a tensor of pseudotime-series GRNs, which is ready for subsequent dynamic network analyses. To facilitate the building of workflows, “modular” functions are developed and included in scGEAToolbox. These modular functions perform basic common tasks that are required by different tools. For example, a modular function uses different methods to compute the cell-to-cell similarity matrix and another modular function uses different methods to estimate the number of cell clusters. In addition to existing methods, several new functions are present in scGEAToolbox. These include a function for feature gene selection using three summary statistics: expression mean (μ), coefficient of variation (CV) and the dropout rate (r_drop_) and a function for trajectory construction based on the splinefit algorithm (Lundgren, 2019). Most functions in scGEAToolbox can be called through the GUI application without using the command line. This feature makes scGEAToolbox a useful training tool for beginners. Visualization functions support intuitive interpretations of the data. The source code of scGEAToolbox is provided free for academic use. When needed, stand-alone applications of scGEApp can be built for all major platforms with or without Matlab installed. In summary, scGEAToolbox provides comprehensive analysis support for scRNA-seq data in Matlab. It makes two key contributions: (1) implementing and incorporating a large number of high-level functions, and (2) defining an easy-to-use GUI for commonly used methods. I anticipate that these key features will make scGEAToolbox a useful tool for researchers to conduct analyses with scRNA-seq data and develop new algorithms more efficiently.

## Acknowledgements

The author thanks Daniel Osorio, Jianhua Huang, Yan Zhong and Guanxun Li for helpful discussion and inspiration during the development of this software tool.

